# Genetic variation of human myokine signaling is dominated by biologic sex and sex hormones

**DOI:** 10.1101/2022.01.20.477045

**Authors:** Leandro M. Velez, Cassandra Van, Timothy M. Moore, Zhenqi Zhou, Casey Johnson, Andrea L. Hevener, Marcus M. Seldin

## Abstract

Proteins secreted from skeletal muscle, termed myokines, allow muscle to impact systemic physiology and disease. Myokines play critical roles in a variety of processes, including metabolic homeostasis, exercise improvements, inflammation, cancer and cognitive functions^1–6^. Despite the clear relevance of these factors in mediating a multitude of physiological outcomes, the genetic architecture, regulation and functions of myokines, as well as degree of conservation of these communication circuits remains inadequately understood. Given that biologic sex controls critical aspects of nearly every physiologic outcome, it is essential to consider when relating specific mechanisms to complex genetic and metabolic interactions. Specifically, many metabolic traits impacted by myokines show striking sex differences arising from hormonal^7–10^, genetic^7,11^ or gene-by-sex interactions^12,13^. In this study, we performed a genetic survey of myokine gene regulation and cross-tissue signaling in humans where sex as a biological variable was emphasized. While expression levels of a majority of myokines and cell proportions within skeletal muscle showed little differences between males and females, nearly all significant cross-tissue enrichments operated in a sex-specific or hormone-dependent fashion; in particular, with estrogens. These sex- and hormone-specific effects were consistent across key metabolic tissues: liver, pancreas, hypothalamus, intestine, heart, visceral and subcutaneous adipose tissue. Skeletal muscle estrogen receptor enrichments across metabolic tissues appeared stronger than androgen receptor and, surprisingly, ~3-fold higher in males compared to females. To define the causal roles of estrogen signaling on myokine gene expression and functions, we generated male and female mice which lack estrogen receptor α (*Esr1*) specifically in skeletal muscle and integrated global RNA-Sequencing with human data. These analyses highlighted mechanisms of sex-dependent myokine signaling conserved between species, such as myostatin enriched for divergent substrate utilization pathways between sexes. Several other sex-dependent mechanisms of myokine signaling were uncovered, such as muscle-derived *TNF*α exerting stronger inflammatory signaling in females compared to males and *GPX3* as a male-specific link between glycolytic fiber abundance and hepatic inflammation. Collectively, we provide the first genetic survey of human myokines and highlight sex and estrogen receptor signaling as critical variables when assaying myokine functions and how changes in cell composition impact other metabolic organs.

## Results

### Sex hormones, but not biologic sex show stronger enrichment with myokine expression

Our goal was to perform a comprehensive survey how skeletal muscle communicates with and impacts metabolic organs. We focused these analyses on exploiting natural genetic variation to assay muscle-specific regulation of myokines and changes in cellular composition, then relate these outcomes to consequent cross-tissue signaling mechanisms (Fig 1A). Initially, we quantified differential expression of genes encoding all known secreted proteins in skeletal muscle from 210 male and 100 female individuals^14^. While several notable myokines appeared different between sexes (Fig 1B), a striking majority of all muscle secreted proteins (74%) showed no difference in expression between males and females (Fig 1C, Supplemental table 1). To understand potential sex-effects on the regulation of myokines, gene ontology enrichments were performed in muscle. Specifically, the skeletal muscle genes which showed the strongest correlation with myokines corresponding to each category (male-specific, female-specific or non-sex specific) were used for pathway enrichments. Here, the top 10 pathways which persisted in females were also observed strongly enriched within the non-sex-specific category, whereas pathways enriched for male-specific myokines were distinct (Fig 1D). Notably, the female and shared pathways suggested roles in epigenetics and RNA processing, while male-specific myokine coregulated processes were more enriched in metabolic pathways (ex. NADH metabolism) (Fig 1D). Further, a majority of myokines showed strong correlation with receptors mediating functions of androgens (androgen receptor -*AR*), estrogens (*ESR1*), or both, regardless of sex-specific expression (Fig 1E). We note that expression of hormone receptors themselves were also not significantly different between sexes (Figure 1 - Figure supplement 1). To infer causality from hormone receptor regulation, we performed RNA-sequencing on mice lacking estrogen receptor α (*Esr1*) in skeletal muscle specifically and integrated these analyses with human myokine estimates. While myokines not regulated by *ESR1* showed little sex-specific modes of expression, those which were estrogen-dependent showed much stronger representation of sex-specificity, in particular in males (Fig 1F-G). Among these, the master regulator of skeletal muscle differentiation and proliferation, myostatin (*MSTN*), was strongly correlated with *ESR1* and *AR* in both sexes. Despite these hormone receptor correlations, the gene was markedly higher in males compared to females, where ablation of *ESR1* uniquely drove expression (Fig 1H). These data suggest interactions between biologic sex and *ESR1* to tightly regulate *MSTN* in males, where other factors could contribute more in females. This sex-specific regulation of myostatin also showed differences in functional annotations, as the most highly enriched pathways in males showed GO terms related to glycolytic metabolism compared to oxidative phosphorylation in females (Fig 1I). These observations are consistent with previous studies which note myostatin-dependent increases in muscle mass in males, but not females^15,16^, where estrogen signaling is suggested as a mechanism mediating these differences. These data demonstrate that, expression of most myokines is not different between biologic sexes; however, interactions between sex and hormone receptors likely play important roles in determining myokine regulation and local signaling.

**Figure 1.**
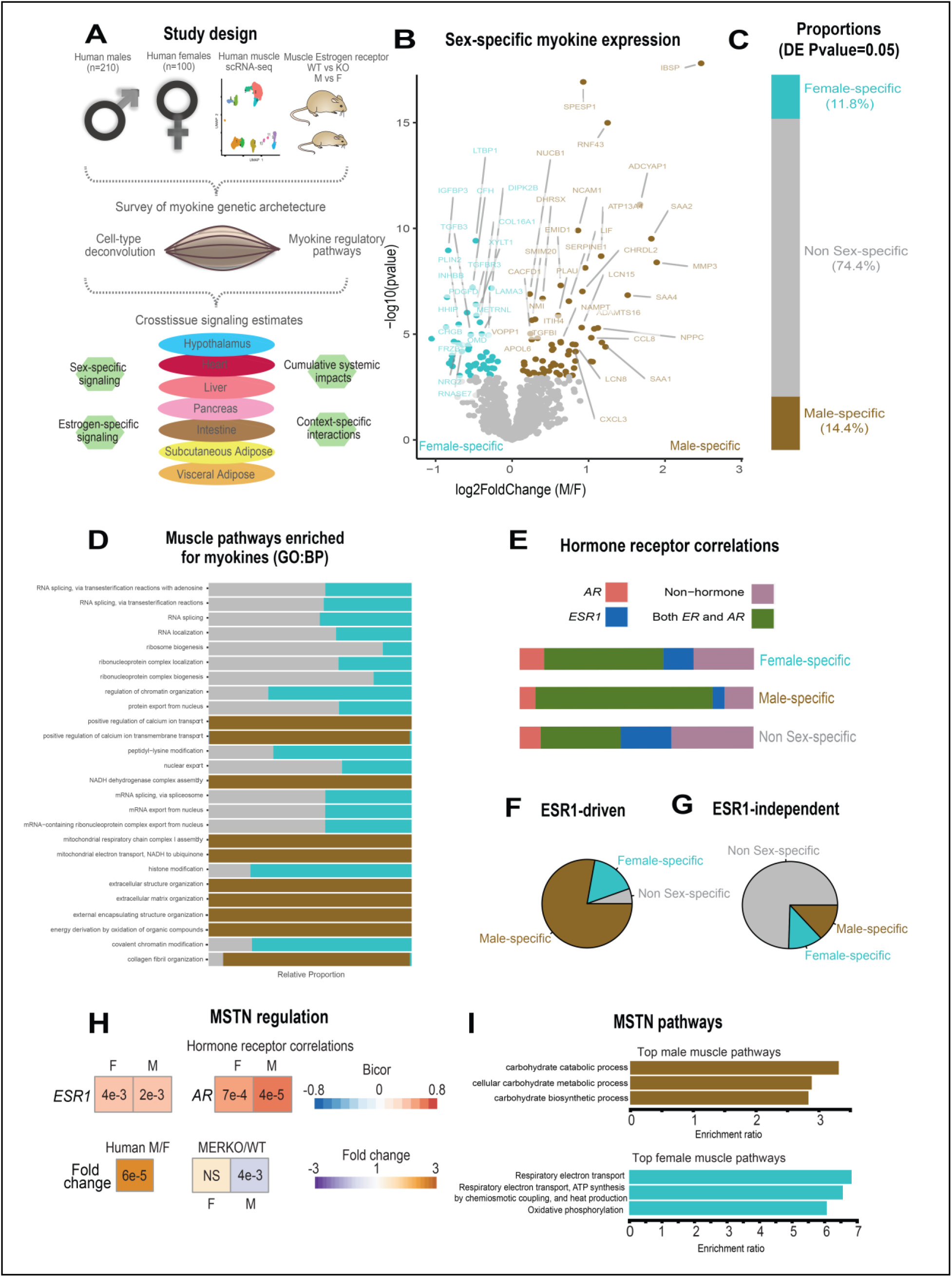
Sex and hormone effects on myokine regulation. A, overall study design for integration of gene expression from muscle from 310 humans, single-cell RNA-seq, muscle-specific deletion of *Esr1* to infer interorgan coregulatory process across major metabolic tissues. B-C, Differential expression analysis for sex was performed on all genes corresponding to secreted proteins in skeletal muscle (Myokines). The specific genes which showed significant changes in each sex are shown as a volcano plot (B) and the relative proportions of myokines corresponding to each category at a modest logistic regression p-value less than 0.05 (C). D, for each differential expression category based on sex shown in C, myokines were correlated with all other muscle genes for pathway enrichment. Then the top 10 enriched pathways in males, females, or non-sex specific (by overall significance) were visualized together where number of genes corresponding to each category shown as a relative proportion. E, the same analysis as in D, except instead of myokines being correlated with all muscle genes, they are binned into proportions correlated with AR, ESR1, both hormone receptors, or neither. F-G, Myokines were binned into 2 categories based on significantly differentially expressed (logistic regression adjusted p-value<0.05) between muscle-specific WT and *Esr1*-KO mice (F) or those that showed no change (G), then visualized as relative proportions within each category shown in C. H, Midweight bicorrelation (bicor) coefficients (color scheme) and corresponding regression p-values (filled text) are shown for muscle MSTN and ESR1 or AR in both sexes (top). Below correlations are shown differential expression log2FC (color scheme) and corresponding logistic regression p-values (text fill) for MSTN between sexes in humans or WT vs muscle-specific ESR1 KO mice (MERKO). I, the top 3 pathways of genes which significantly (p<1e-4) correlated with muscle MSTN in males (top) or females (bottom). For human data, n=210 males and n=100 females. For mouse MERKO vs WT comparisons, n=3mice per group per sex. p-values from midweight bicorrelations were calculated using the students p-value from WGCNA and logistic regression p-values were calculated using DESeq2.

### Sex dominates cross-tissue pathways enriched for myokines

Given that expression levels of most myokines appeared similar between sexes, we next assessed putative functions across organs. We applied a statistical method developed to infer cross-tissue signaling which occur as a result of genetic variation^17–19^. Here, we assayed the distribution of midweight bicorrelation coefficients between myokine expression levels and global gene expression measures from the same individuals in key metabolic tissues including hypothalamus, heart, intestine, pancreas, liver, subcutaneous and visceral adipose tissue. Remarkably, nearly all highly significant correlations between myokines and target organ genes showed sex-specific modes of operation (Fig 2A-H). This sex specificity also appeared more pronounced for positive correlations between myokines and target tissue genes, as compared to negative (Fig 2-H). Further, among these high significant cross-tissue circuits, myokine hormone receptor enrichment was strongly dependent on the category (ex. significant only in females) rather than target tissue (Fig 2A-H). This observation further suggests that hormone receptor levels (*ESR1* or *AR*) in muscle are a stronger determinant of myokine expression compared to biologic sex; however, sex is suggested to dominate coregulated signaling processes across organs via myokines. Therefore, to gauge the relative impact of muscle steroid hormone receptors across organs, the number of significant correlations between *ESR1*, *AR* or both were quantified for each tissue. Remarkably, *ESR1* specifically as showing an order of magnitude stronger enrichment across metabolic tissues in compared to *AR* or both, where the number of significantly correlated cross-tissue male *ESR1* genes (Fig 2I) were three-fold higher than females (Fig 2J). Because both sex and *ESR1* signaling appeared critical in the regulation of myokine functions, we binned significant cross-tissue enrichments into categories taking into consideration whether myokines were driven by *ESR1* in muscle, and/or showing a sex-specific mode of inter-organ enrichment. This analysis suggested that a majority of myokines were either driven by *ESR1* and signaled robustly across sexes (Fig 2K, yellow) or signaled differently between sexes, but regulated independent of *ESR1* (Fig 2K, red). These categories appeared to a much greater result as opposed to a combination of both ESR1-driven myokine and sex-specific cross-tissue signaling (Fig 2K, beige) or neither (Fig 2K, seagreen). One notable example of sex-specific signaling was observed for tumor necrosis factor alpha (TNFα) which showed markedly different putative target tissues (Fig 2L, left), as well as underlying functional pathways (Fig 2L, right), depending on sex. For example, overall inflammatory processes engaged by TNFα were substantially stronger in adipose tissue in females; however, the same pathways were higher in liver and hypothalamus in males (Fig 2L, left). Collectively, these data show that sex and related sex steroid hormones, particularly estradiol, exert dominant roles in regulating tissue and pathway engagement by myokines.

**Figure 2.**
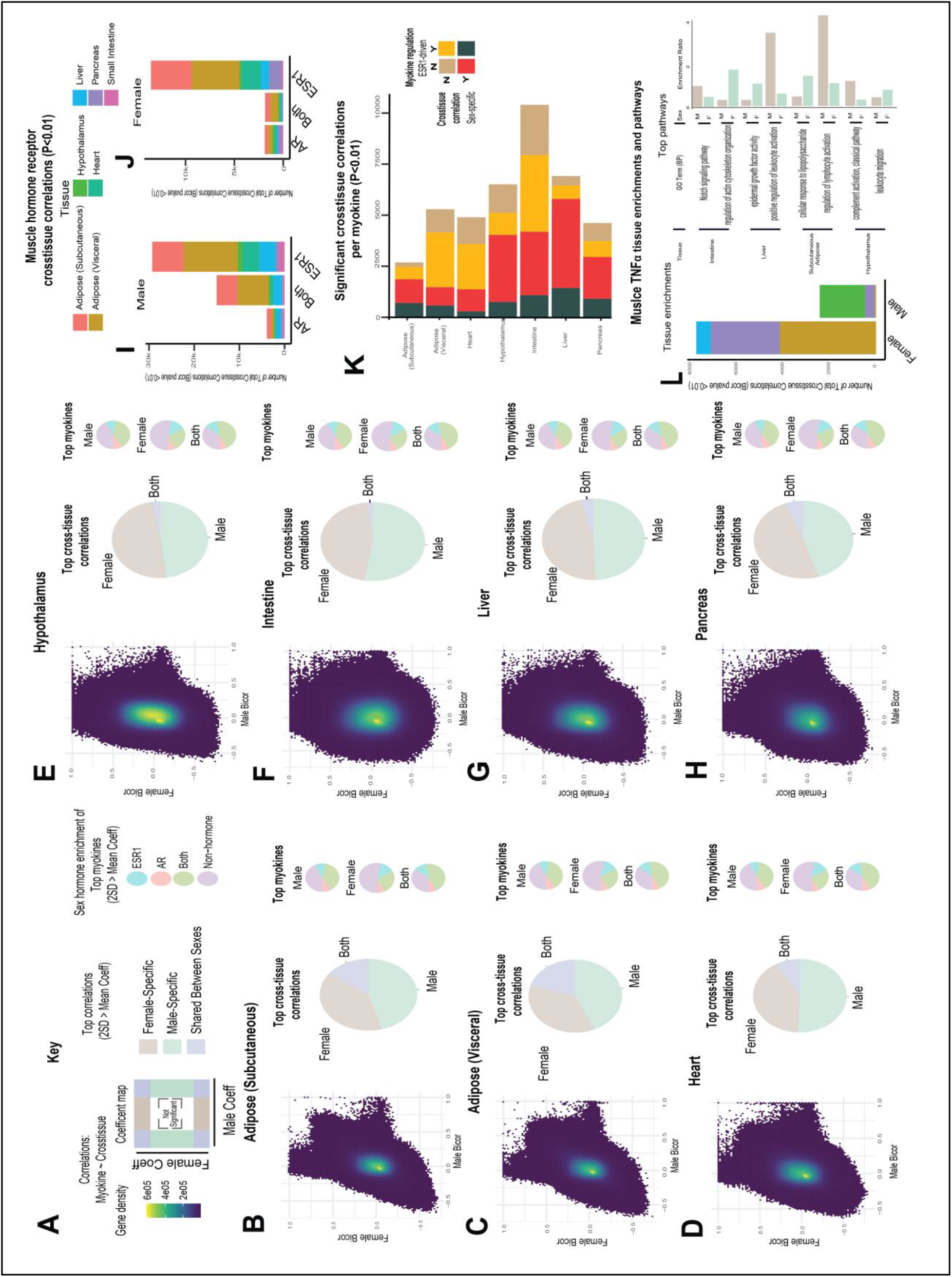
Sex and hormone effects on myokine regulation. A-H, Key illustrating analysis for distribution of midweight bicorrelation coefficients between all myokines in skeletal muscle and global transcriptome measures are plotted between sexes (left), where proportions for 2SD > mean are subdivided into occurrence uniquely in females, males, or shared (middle). The myokines identified in each category were then binned into hormone receptor correlations for *ESR1, AR*, both or neither (right). This analysis was performed on all myokines across subcutaneous adipose tissue (B), visceral adipose (C), heart (D), hypothalamus (E), small intestine (F), liver (G) and pancreas (H). I-J, Significant cross-tissue correlations between muscle *ESR1, AR*, or both hormone receptors are colored by tissue and shown for males (I) or females (J). K, For each tissue (y-axis), the ratio of significant cross-tissue correlations per muscle myokine (x-axis) are shown and colored by categories of: either the myokine regulated by ESR1 and/or the target tissue regression occurring specifically in one sex. L, Number of significant cross-tissue correlations with muscle TNFα are shown for each sex and colored by tissue as in I-L (left). The −log10(p-value) of significance in an overrepresentation test (x-axis) are shown for top significant intertissue pathways for muscle TNFα in each sex (right).

### Muscle cell proportions are similar between sexes, but associated changes across tissues show sex-specificity

To determine the impact of muscle composition on other tissues, we next surveyed muscle cellular proportions in the context of genetics and sex. Here, we integrated single-cell sequencing of human skeletal muscle^20^ using cellular deconvolution^21^ to estimate cellular composition (Fig 3A). Here, a proportions in admixture approach^22^ outperformed other methods (Figure 3 - Figure supplement 1) to capture a majority of established cell populations across individuals (Supplemental Table 2). Similar to myokine expression, no differences were observed between sexes in terms of cell composition, with the exception of modest higher glycolytic fiber in males, compared to elevated oxidative fiber levels in females (Fig 3B). Additionally, no differences were observed in the correlations between compositions within males or females (Figure 3 - Figure supplement 2); however, nearly every cross-tissue enrichment corresponding to an individual muscle cell type differed between sexes (Fig 3C). Generally, differences in skeletal muscle cell abundance corresponded to changes in liver and visceral adipose tissue pathways in males, compared to pancreas in females (Fig 3C). In contrast to general myokine enrichments, cell proportions showed stronger correlations with *AR* when compared to *ESR1* across both sexes; however, the most abundant cell types were significantly enriched for both steroid hormone receptors (Fig 3D). Next, to uncover potential direct mechanisms linking changes in cell composition to peripheral tissues, we analyzed associated myokines and adopted and adjusted regression-based mediation approach. Despite few differences between sexes in terms of myokine expression and cell composition, specific myokines highly correlating with individual cell type were markedly different between males and females with the exception of one, APOD in slow-twitch fibers (Fig 3E). To determine if variation in cell compositions corresponding to sex-specific tissue signaling via myokines was predicted to be causal, we implemented adjusted regression mediation analyses^23,24^ for glycolytic fiber composition. Because male glycolytic fiber type was selectively enriched for liver pathways such as immune cell activation and regulated exocytosis (Fig 3F), the top-genes driving these enrichments were used to determine causality. The top-correlated muscle secreted protein with male glycolytic fiber type levels was secreted glutathione peroxidase 3 (*GPX3*). Here, adjusting regressions between glycolytic fiber and liver pathways on *GPX3* significantly reduced the overall significance across tissues (Fig 3G), suggesting *GPX3* as a mediator of this communication. These data point to a potential mechanism whereby muscle fiber abundance could buffer free radical generation in the liver, thereby feeding back on inflammation. This analysis appeared additionally sensitive to inferring non-dependent relationships between muscle cell types, top-ranked myokines and cross-tissue processes. For example, female glycolytic fibers were strongly enriched for pancreatic protein synthesis pathways; however, when adjusted for the top-ranked myokine *CES4A*, no changes in regression significance were observed (Fig 3F-G). These analyses show that male *GPX3* is a likely mechanism whereby fast-twitch muscle signals to liver; however, the same cell type in females drive pancreas protein synthesis independent of *CES4A*. In summary, we show that cell composition is strongly conserved between sexes, but cross-tissue signaling of altered composition differs entirely. We further suggest putative myokines and mechanisms, as well as highlight the key regulatory roles of estrogen in both sexes.

**Figure 3.**
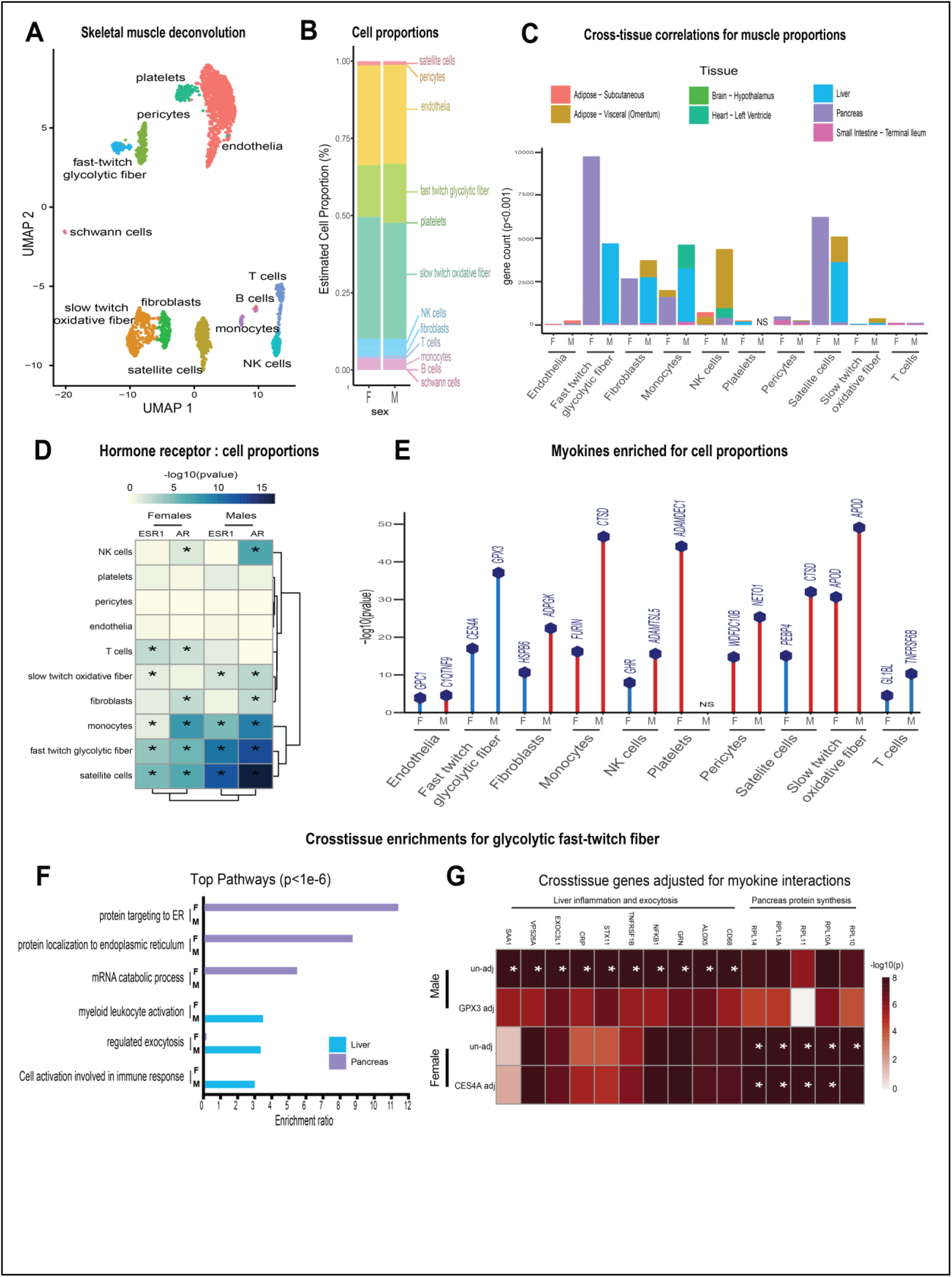
Genetic variation of muscle cell proportions and coregulated cross-tissue processes. A, Uniform Manifold Approximation and Projection (UMAP) for skeletal muscle single-cell sequencing to deconvolute proportions. B, Mean relative proportions of pseudo-single-cell muscle cell compositions (denoted by color) between sexes. C, Number of significant cross-tissue correlations (y-axis) corresponding to each skeletal muscle type in each sex (x-axis). Target tissues are distinguished by color, where NS (male platelets) denotes that no significant cross-tissue correlations were observed. D, Heatmap showing significance of correlations between skeletal muscle hormone receptors and cell proportions, * = p<0.01. E, the strongest enriched myokines are plotted for each myokine (y-axis, −log10p-value of myokine ~ cell composition) are shown for each muscle proportion for each sex (x-axis). Gene symbols for myokines are shown above each line, where red lines indicate positive correlations between myokine and cell type and blue shows inverse relationships. F, Significant cross-tissue correlated genes in liver (blue) and pancreas (purple) for muscle fast twitch glycolytic fibers (P<1e-6) were used for overrepresentation tests where enrichment ratio of significance (x-axis) is shown for each pathway and sex (y-axis). G, Heatmap showing the regression significance of the top 5 genes corresponding to inflammation (liver), exocytosis (liver) and protein synthesis (pancreas) for proportions of fast-twitch fiber type (un-adj). Below each correlation between fast-twitch fiber and liver or pancreas gene, the same regressions were performed while adjusting for abundance of select myokines in each sex. *=p<1e-6.

### Conclusions and limitations

Here we provide a population survey of skeletal muscle myokine regulation and putative functions using genetic variation and multi-tissue gene expression data. We find that in general, expression of myokines do not significantly differ between sexes; however, signaling mechanisms across tissues inferred from regressions show strong sex specificity. Steroid hormones, in particular *ESR1*, is highlighted as a key regulator of myokines and potentially interacting with biologic sex for proteins such as myostatin. Further integration with loss-of-function mouse models of *Esr1* highlighted the key roles of estradiol signaling in muscle in terms of myokine regulation and signaling across both sexes. Generation of pseudo-single-cell maps of muscle composition showed that, like myokines, muscle proportions are conserved between sexes, but inferred interorgan consequences differ substantially. When interpreting these findings, serval considerations should be taken. While inter-tissue regression analyses have been informative to dissect mechanisms of endocrinology^17–19,25^, observations can be subjected to spurious or latent relationships in the data. While causality for inter-organ signaling can be inferred statistically using approaches such as mediation as in Fig 3H, the only methods to provide definitive validation for new mechanisms is in experimental settings. In addition, we anticipate that estimates for *ESR1* effects on myokines in this study likely represents an underestimated number of all human *ESR1*-driven myokines. The primary limitation here is that annotation of known orthologous mouse-human genes^26^ is limited to roughly ~15% of the coding genome. Related, integration with muscle specific AR-deletion would offer intriguing information for cell composition metrics, but also limited similarly. Furthermore, cell composition estimates from single-cell sequencing data are inferred from gene expression, where histological or flow cytometry-based methods can provide more accurate direct quantifications. Clearly, morphological and structural differences between sexes have been observed in humans^27^ which, if not apparent in gene expression, would be missed in this analysis. Future studies addressing these points will help to clarify context- and mechanism-relevant muscle-derived endocrine communication axes. In summary, this study highlights the critical nature of sex and sex steroid hormones in mediating myokine functions which should be considered when interpreting future studies of myokines.

## Material and methods

### All datasets used, R scripts implemented for analyses and detailed walkthrough guide is available via:https://github.com/marcus-seldin/myokine-signaling

#### Data sources and availability

All data used in this study can be immediately accessed via github to facilitate analysis. Human skeletal muscle and metabolic tissue data was accessed through GTEx V8 downloads portal on August 18, 2021 and previously described^14^. To enable sufficient integration and cross-tissue analyses, these data were filtered to retain genes which were detected across tissues where individuals were required to show counts > 0 in 1.2e6 gene-tissue combinations across all data. Given that our goal was to look across tissues at enrichments, this was done to limit spurious influence of genes only expressed in specific tissues in specific individuals. Post-filtering consists of 310 individuals and 1.8e7 gene-tissue combinations). Single-cell sequencing from skeletal muscle used for deconvolution was obtained from^20^. *Esr1* WT and KO mouse differential expression results are available on Github as well, where raw sequencing data has been deposited in NIH sequence read archive (SRA) under the project accession: PRJNA785746

#### Selection of secreted proteins

To determine which genes encode proteins known to be secreted as myokines, gene lists were accessed from the Universal Protein Resource which has compiled literature annotations terms for secretion^28^. Specifically, the query terms to access these lists were: locations:(location:“Secreted [SL-0243]” type:component) AND organism:“Homo sapiens (Human) [9606]” where 3666 total entries were found.

#### Differential expression of myokines dependent on sex

Gene expression counts matrices were isolated from the rest of the tissues, where individual genes were retained if the total number of counts exceeded 10 in 50 individuals. Next, only genes encoding secreted proteins (above) were retained, where logistic regression contrasted on sex was performed using DESeq2. Differential expression summary statistics were used for downstream binning of sex-specificity based on an empirical logistic regression pvalue <0.05. This threshold was used to reflect a least stringent cutoff where, despite potential false positive influence, genes which nominally trended toward sex-specific expression could be included in those categories. Given that the general conclusions supported very few proportions of myokines showing sex-specific patterns of expression, this conclusion would only be further exaggerated if the DE threshold were made more stringent and lessened the number of myokines in each category.

#### Regression analyses across tissues

Regression coefficients and corresponding p-values across tissues were generated using WGCNA bicorandpvalue() function^22^. Myokine-target gene pairs were considered significant (ex. Fig 2A-H) at a threshold of abs(bicor) > 2 standard deviations beyond the average coefficient for the given target tissue of interest. In previous studies, this threshold of 2 standard deviations reflects adaptive permutation testing pvalues <0.01^17,18^. For analyses estimating cumulative patterns of concordance across tissues (ex. Fig 2I-L), empirical regression pvalues (students pvalue from bicor coefficients) of 0.01 (corresponding to abs(bicor)>0.1) were used to assay global patterns. While empirical pvalues are subjected to false positives, including these enables broad visualization of both potential direct interactions (ex. myokine-target gene) as well as coregulated processes across organs. It is important to note that we exclusively rely on these empirical pvalues when surveying broad correlation structures, whereas much more stringent and appropriate thresholds (ex. p<1e-6 for Fig 3G) were applied when inferring direct interactions.

#### Pathway enrichment analyses

For Fig 1I and Fig 3G, genes corresponding to pvalue cutoffs were visualized using Webgestalt^29^ to enable streamline visualization. For Fig. 1D, the top 1000 (by regression p-value) significant genes from myokines to all muscle bicorrelation analysis in females, males or non-sex specific datasets were assessed for enrichment in GO Biological Process terms using ClusterProfiler ver. 4.0.2 in R^30^. The resulting top ten GO terms in each dataset were integrated and plotted against the relative proportion of the p.adjusted value, and visualized in the same graph using ggplot2.

#### Deconvolution of skeletal muscle

Raw single-cell RNA sequencing from skeletal muscle was obtained from^20^. These raw counts were analyzed in Seurat where cluster analyses identified variable cell compositions. Cell type annotations were assigned based on the top 30 genes (Supplemental table 2) assigned to each UMAP cluster through manual inspection and ENRICHR^31^. Finally, a normalized matrix of gene:cells was exported from Seurat and used to run deconvolution on skeletal muscle bulk sequencing. Using the ADAPTS pipeline^21^, three deconvolution methods (nnls, dcq or proportions in admixture) were compared based on ability to robustly capture cell proportions (Supplemental fig 2), where proportion in admixture showed the best performance and subsequently applied to bulk sequencing.

#### ESR1 muscle KO generation, RNA-Seq and integration with human data

Muscle-specific Esr1 deletion was generated and characterized as previously described^10^. Whole quadriceps was pulverized at the temperature of liquid nitrogen. Tissue was homogenized in Trizol (Invitrogen, Carlsbad, CA, USA), RNA was isolated using the RNeasy Isolation Kit (Qiagen, Hilden, Germany), and then tested for concentration and quality with samples where RIN > 7.0 used in downstream applications. Libraries were prepared using KAPA mRNA HyperPrep Kits and KAPA Dual Index Adapters (Roche, Basel, Switzerland) per manufacturer’s instructions. A total of 800-1000 ng of RNA was used for library preparation with settings 200-300 bp and 12 PCR cycles. The resultant libraries were tested for quality. Individual libraries were pooled and sequenced using a HiSeq 3000 at the UCLA Technology Center for Genomics and Bioinformatics (TCGB) following in house established protocols. Raw RNAseq reads were inspected for quality using FastQC v0.11.9 (Barbraham Institute, Barbraham, England). Reads were aligned and counted using the Rsubread v2.0.0^32^ package in R v3.6 against the Ensembl mouse transcriptome (v97) to obtain counts. Lowly expressed genes (>80% samples with 0 counts for particular gene) were removed. Samples were analyzed for differential expression using DeSeq2 v1.28.0^33^.

#### Conservation of gene between mice and humans

To find which myokines and pathways were conserved between mice and humans, all orthologous genes were accessed from MGI vertebrate homology datasets, which have been compiled from the Alliance for Genome Resources^26^ and intersected at the gene level.

## Acknowledgements

We acknowledge the following funding sources for supporting these studies: LMV, CV, CJ and MMS were supported by NIH grants HL138193, DK130640 and DK097771. ZZ was supported by NIH grant DK125354. T.M.M. was supported by the UCLA Intercampus Medical Genetics Training Program (T32GM008243). ALH is supported by NIH grants U54 DK120342, R01 DK109724, and P30 DK063491.

## Author contributions

LV, CV, TMM, ZQ and MMS accessed raw data, performed analyses and drafted the manuscript. CJ and AH provided critical insight into data use and interpretation, as well as guided the study. All authors read and approved this manuscript.

---

**Figure 1 - Figure supplement 1:**
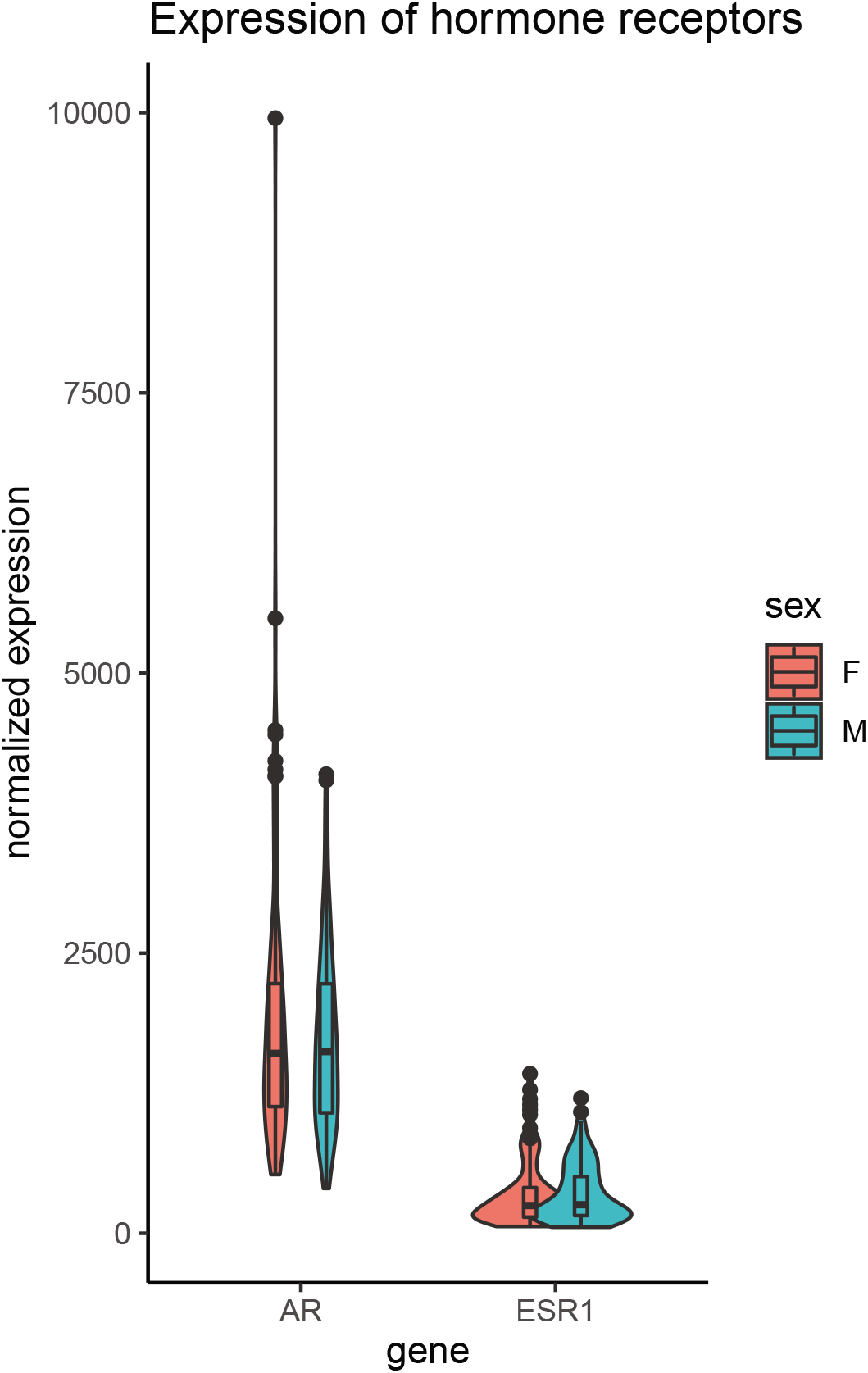
Skeletal muscle sex hormone receptor expression between sexes: Normalized gene expression levels for androgen receptor (AR) or estrogen receptor (ESR1) (y-axis) in each sex (x-axis). None of the expression levels were significantly different between sexes (students t-test, two-way)

**Figure 3 - Figure supplement 1:**
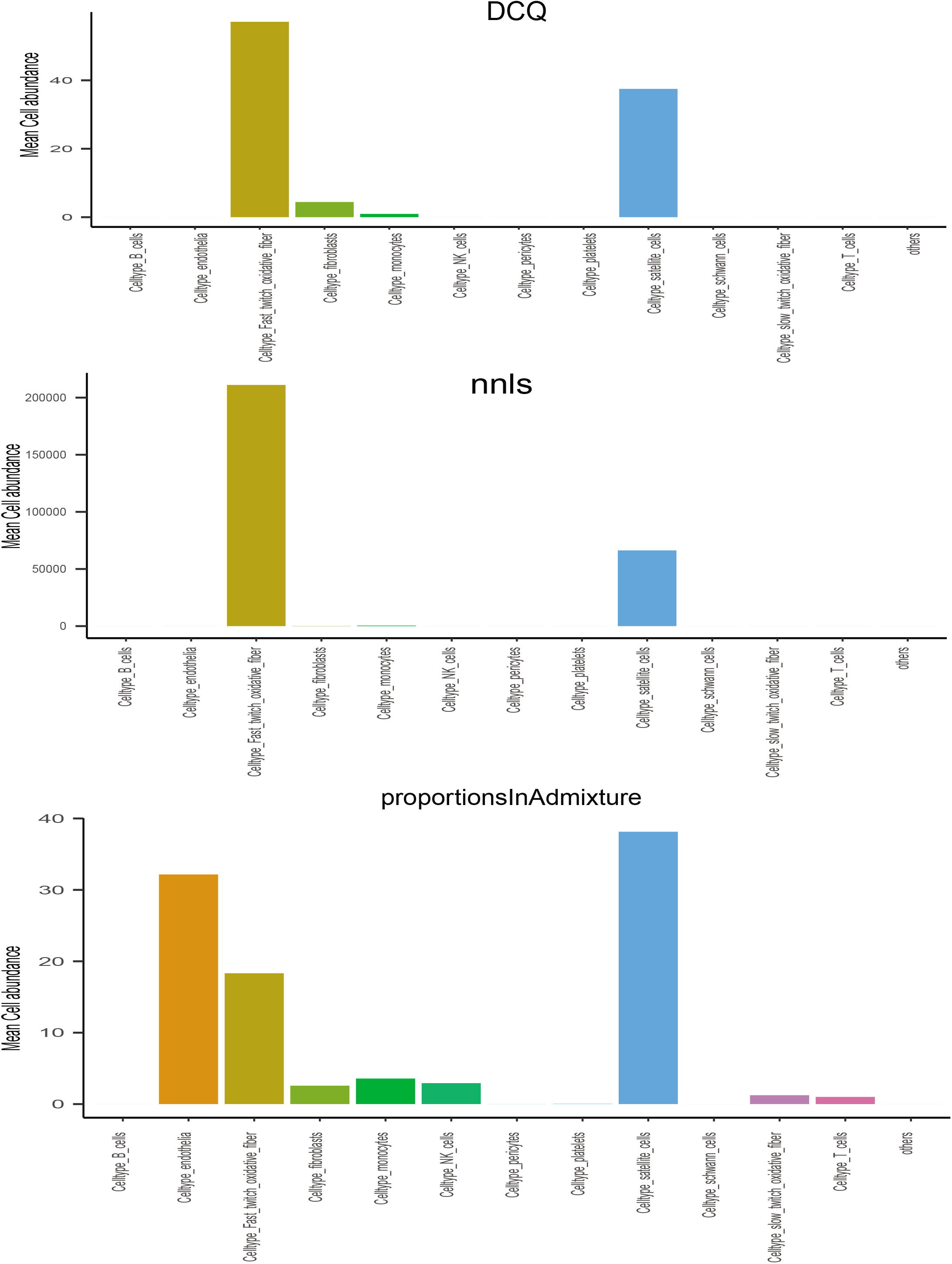
Comparisons of deconvolution methods. Cell proportions were estimated from skeletal muscle sequncing across the 310 individuals in GTEx. Here, comparisons of the three most common methods (DCQ, NNLS and porportionsInAdmixture) were plotted for each pseudo-sc-proportion, where proportionsInAdmixture method captured the largest relative number of cell types

**Figure 3 - Figure supplement 2:**
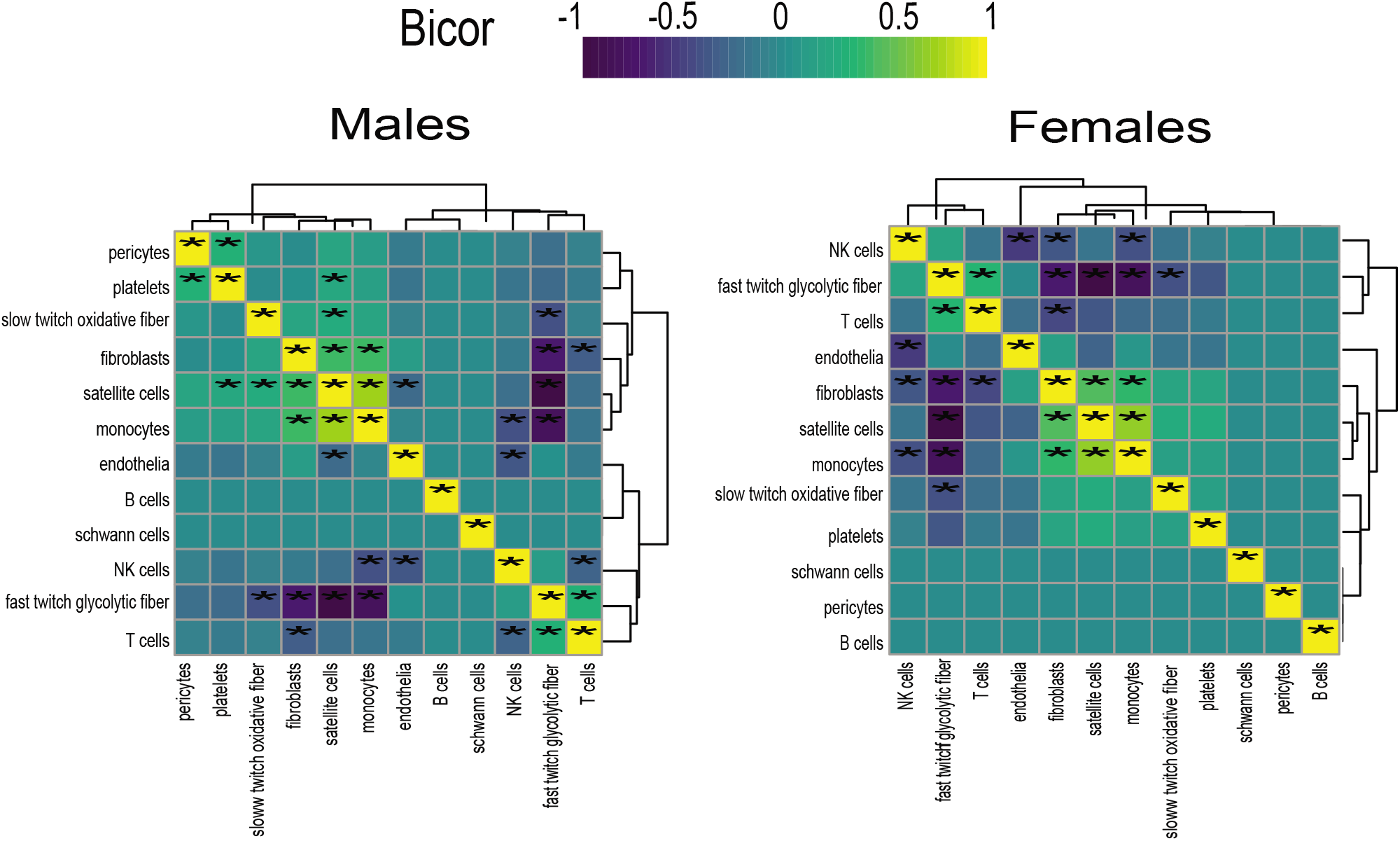
Cell composition correlations within each sex: Heatmaps showing regressions for cell proportions in males (left) or females (right), * = regression pvalue<0.01

